# Inheritance of OCT4 predetermines fate choice in human embryonic stem cells

**DOI:** 10.1101/137299

**Authors:** Samuel C. Wolff, Raluca Dumitru, Katarzyna M. Kedziora, Cierra D. Dungee, Tarek M. Zikry, Rachel A. Haggerty, JrGang Cheng, Adriana S. Beltran, Jeremy E. Purvis

**Author notes:** Corresponding Author: Jeremy Purvis, Genetic Medicine Building 5061, CB#7264, 120 Mason Farm Road, Chapel Hill, NC 27599-7264.

## Abstract

Clonal cells can make different fate decisions, but it is often unclear to what extent these decisions are autonomous or predetermined. Here, we introduce a live-cell reporter for the pluripotency factor OCT4 into human embryonic stem cells to understand how they choose between self-renewal and differentiation. By tracing the histories of individual cells over multiple generations, we found that cells whose offspring were destined to differentiate showed decreased expression of OCT4 long before exposure to the differentiation stimulus. OCT4 levels were lineage-dependent; however, during cell division, mother cells distributed OCT4 asymmetrically to daughters. The resulting ratio of OCT4 between sister cells—established within minutes of mitosis—was predictive of downstream fates: cells receiving a greater ratio of maternal OCT4 showed sustained OCT4 levels and a reduced capacity to differentiate. Our observations imply that the choice between two developmental fates is almost entirely predetermined at the time of cell birth through inheritance of a pluripotency factor.

## Introduction

It is well established that clonal cells can make distinctly different fate decisions (Suda et al., 1984a; Suda et al., 1984b; Suel et al., 2006). An important conceptual challenge, however, is to understand to what extent a cell exerts independent control over its own fate (Symmons and Raj, 2016). At one extreme, a cell’s fate may be entirely shaped through environmental stimuli and autonomous signaling; at the other extreme, its fate already determined before it emerges from its mother cell, rendering it impervious to external cues. Although multiple studies have shown that different single-cell signaling patterns are associated with different downstream responses (Albeck et al., 2013; Lane et al., 2017; Lin et al., 2015; Purvis et al., 2012), tracking cells over multiple cell-cycle generations suggests that intracellular signals themselves can be inherited from mother to daughter cells. For example, the response to death ligands in human cells is similar among sister cells through inheritance of apoptotic protein factors (Spencer et al., 2009). In addition, cell-cycle checkpoint decisions in daughter cells were shown to be influenced by the signaling history of the mother cell (Arora et al., 2017; Barr et al., 2017; Yang et al., 2017). Taken together, these studies have eroded the concept that cells can make fully autonomous fate decisions and raise the question of what mechanisms may mediate the inheritance of fate-determining factors.

How fate choice is controlled is also a central question in stem cell biology. During human development, proliferating stem cells give rise to complex and heterogeneous tissues through a dynamic interplay of intracellular signaling events, cell-cell communication, cell proliferation, and morphogen gradients (Deglincerti et al., 2016; Etoc et al., 2016). *In vitro* imaging of attached human embryos has yielded unprecedented insights into the single-cell patterning of the human gastrula (Deglincerti et al., 2016). However, because these cells must be necessarily fixed in preparation for imaging, it is not possible to follow any given cell over the course of its cellular lifetime. It is therefore difficult to pinpoint precisely when an individual cell makes the decision to differentiate, how fate-determining factors are inherited from mother to daughter cell, or why two closely related cells choose different fates. As an alternative approach, human embryonic stem cells (hESCs) represent a promising system for studying embryonic cell fate decisions in real time (Bernardo et al., 2011; Nemashkalo et al., 2017; Thomson et al., 1998). hESCs can be maintained indefinitely in cell culture and are amenable to introduction of fluorescent biosensors to report on intracellular signaling activity (Nemashkalo et al., 2017). These features make it possible for hESCs to be used to understand how fate choice is determined among individual cells.

In this study, we developed an endogenous fluorescent reporter for the human pluripotency factor OCT4 to study its inheritance over multiple cell-cycle generations. We conducted time-lapse fluorescence imaging of hESCs during differentiation to extraembryonic mesoderm and followed the signaling behaviors of individual cells until their final fate decisions were determined. We found that the decision to differentiate is largely determined before the differentiation stimulus is presented to cells and can be predicted by a cell’s preexisting OCT4 levels, bursting frequency, and cell cycle duration. Further, we found that OCT4 levels were highly heritable from mother to daughter cell, but that asymmetric distribution of OCT4 between daughter cell pairs led to sustained differences in OCT4 as well as differences final fate choice. Specifically, differences in OCT4 established within the first hour of cell birth were strongly predictive of cell fate. These results suggest that, in this experimental context, the choice between two developmental fates is nearly predetermined within a short time after stem cell birth.

## Results and Discussion

We first established an experimental system for generating both self-renewing and differentiating cells in response to the same developmental signal (**Figure 1A**). When treated with bone morphogenetic protein 4 (BMP4) for 24 h, a subpopulation of hESCs showed reduced expression of the core pluripotency factor OCT4 and accumulation of the caudal type homeobox 2 (CDX2) transcription factor (**Figure 1B**). Quantitative immunofluorescence (IF) revealed two distinct populations of cells: a pluripotent population with low CDX2 expression that retained the ability to differentiate into other cell types (**Figure EV1**); and a differentiating population of cells with reduced OCT4 expression, increased CDX2 expression, and enlarged morphology (**Figures 1C**). Although BMP4 treatment was originally reported to initiate differentiation toward the trophoblast lineage (Xu et al., 2002), further study has revealed that, in the presence of fibroblast growth factor, it induces markers such as *BRACHYURY* and *ISL1* that are more closely associated with extraembryonic mesoderm (Bernardo et al., 2011). We confirmed the expression of mesodermal markers in BMP4-treated hESCs through quantitative PCR (**Figure EV2**). Thus, treatment of hESCs with BMP4 triggered a binary cell fate decision within 24 hours.

**Figure 1.**
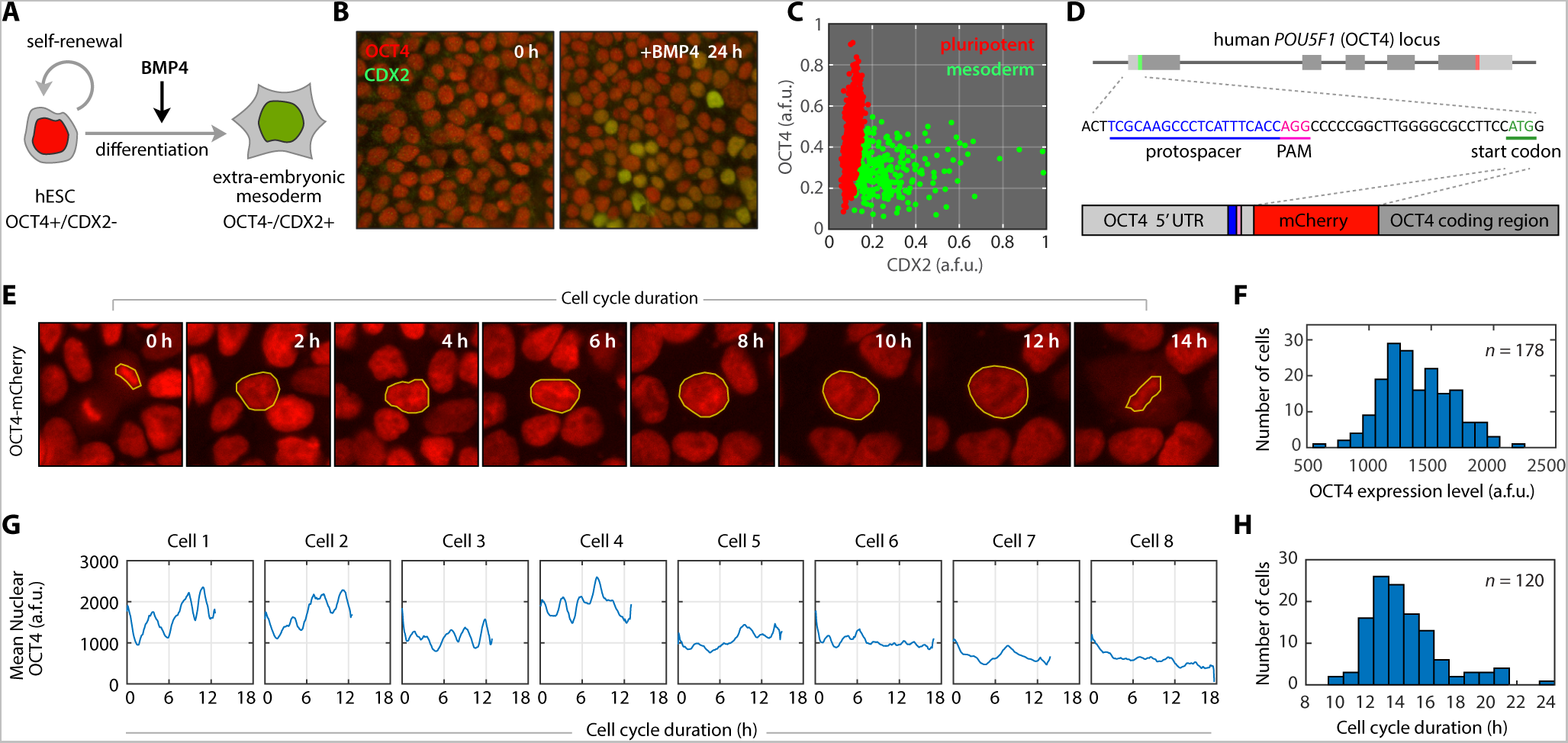
Single-cell dynamics of OCT4 in human embryonic stem cells. **A.** Individual hESCs have the potential to generate another stem cell through self-renewal or to differentiate into a more lineage-specific cell type. **B.** Before differentiation, hESCs show uniformly high expression of OCT4. Treatment with BMP4 produces a mixture of OCT4+/CDX2-self-renewing hESCs and OCT4-/CDX2+ mesodermal cells. **C.** Quantification of OCT4 and CDX2 expression by immunofluorescence after 24 h of BMP4 treatment reveals two populations of hESCs. Cells were assigned to one of two distinct populations based on a 2-component mixed Gaussian distribution (see **Figure EV6**). **D.** A fluorescent mCherry coding sequence was introduced into the endogenous OCT4 locus of H9 hESCs using CRISPR-mediated homologous recombination. **E.** Filmstrip of OCT4 dynamics in an undifferentiated hESC throughout its cell cycle duration. Yellow outlines indicate the region used to quantify mean nuclear fluorescence intensity. **F.** Distribution of OCT4 levels in individual hESCs. A single OCT4 level was quantified for each cell by averaging the mean nuclear mCherry intensity over the lifetime of the cell. **G.** Single-cell traces of OCT4 signaling. The length of each cell’s trace indicates its cell cycle duration. **H.** Distribution of cell cycle durations for 120 hESCs.

To understand how and when individual hESCs make this decision, we developed a fluorescent reporter system to monitor expression of the endogenous OCT4 protein, a canonical marker of the pluripotent state (Nichols et al., 1998). We used CRISPR-mediated genome editing to fuse a monomeric red fluorescent protein (mCherry) to the endogenous OCT4 protein in WA09 (H9) hESCs and isolated a clonal population of single-allele knock-in reporter cells (**Figure 1D** and **Materials and Methods**). The OCT4-mCherry fusion protein showed correct genomic targeting; accurate co-localization with the endogenous OCT4 protein; similar degradation kinetics; and the same chromatin binding pattern near the promoters of OCT4 target genes (**Figure EV3**). Moreover, cells bearing the OCT4-mCherry reporter were competent to differentiate into multiple differentiated cell types (**Figure EV4**), and time-lapse imaging did not alter their proliferation characteristics (**Figure EV5**). For each cell, we calculated a single OCT4 expression level by averaging OCT4-mCherry intensity over its cell cycle duration (**Figure 1E-F**). In addition, we examined the time-series profile of OCT4 dynamics for individual cells and found that the majority of hESCs (68%) displayed sporadic bursts of OCT4 expression that lasted ~1.5 h, with some cells showing as many as 7 bursts (**Figure 1G**). Finally, we calculated individual cell cycle durations, which ranged from 10-24 h with a mean duration of 14.6 h (**Figure 1H**), consistent with the reported population doubling time of ~16 h (Ghule et al., 2011). Thus, our reporter system enabled the reliable analysis of single-cell OCT4 dynamics in hESCs and revealed considerable heterogeneity in untreated stem cells.

With this system in place, we set out to capture the fate decisions of hESCs in real time. First, we performed time-lapse fluorescence imaging of H9 OCT4-mCherry hESCs for 42 h under basal conditions (**Figure 2A**). We then treated these cells with 100 ng/mL BMP4 to induce differentiation while continuing to monitor their responses. Within 12 h of treatment, each cell began to follow one of two distinct fate paths: sustained accumulation of OCT4; or a precipitous decrease in OCT4. After 24 h, cells were fixed and stained for expression of CDX2 to determine their final differentiation status (**Figure 2B**). We imposed a strict cutoff to classify each cell as either self-renewing or differentiated based on its OCT4 and CDX2 expression levels. By fitting the data in **Figure 2B** to a 2-component Gaussian distribution (**Figure EV6**), we selected only those cells that belonged exclusively to either the self-renewing distribution (*p*_*self*_ < 0.01) or the differentiated distribution (*p*_*diff*_ > 0.99), where *p*_*diff*_ represents the probability that a given cell has differentiated. We then traced both populations back through time—spanning multiple cell division events—and labeled each earlier cell according to its “pro-fate”—the fate to which it (or its progeny) would ultimately give rise. The majority of cells in the tracked population were either pro-self-renewing (71%, red traces in **Figure 2A**), giving rise to only self-renewing cells; or pro-differentiated (24%, green traces in **Figure 2A**), giving rise to only differentiated cells. Approximately 5% of cells were “pro-mixed” and gave rise to both fates (yellow traces in **Figure 2A**). Overall, 89% of sister cells chose the same fate, suggesting a large degree of heritability in cell fate and the absence of classically described “asymmetric” cell divisions (Morrison and Kimble, 2006). Thus, time-lapse imaging allowed us to group hESCs by their eventual fate categories before they had received a differentiation signal or had made a clear fate decision. Although the majority of progenitor cells gave rise exclusively to a single fate, a small but significant group of cells gave rise to two different fates.

**Figure 2.**
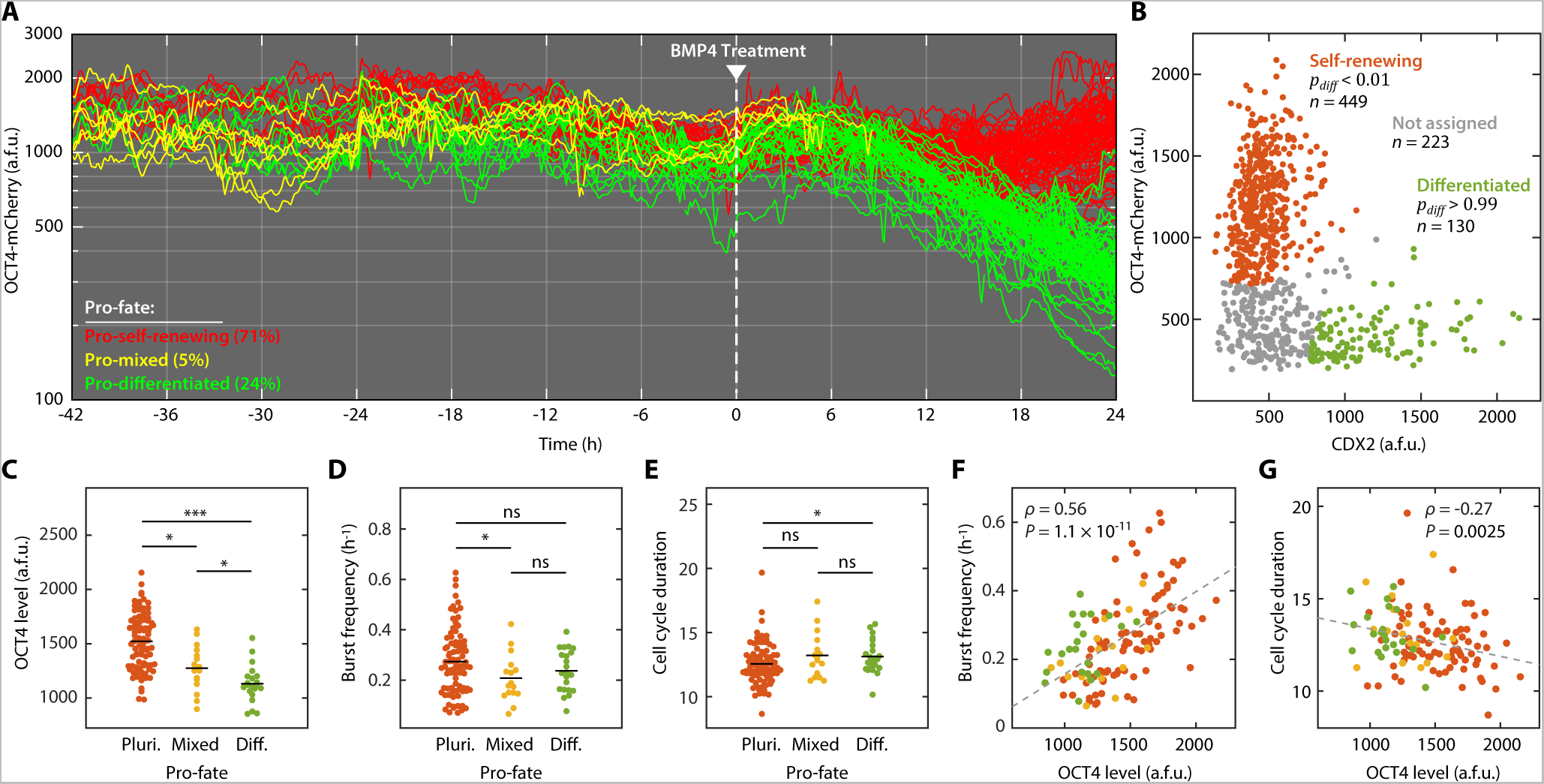
Preexisting differences in OCT4 dynamics predict eventual fate decisions. **A.** Single-cell traces of hESCs before and after treatment with 100 ng/mL BMP4. Cells were imaged for 42 h prior to BMP4 treatment. Mean nuclear OCT4 levels were quantified every 5 minutes and individual cells were tracked from the cell division event that created the cell until its own division. **B.** 24 h after BMP4 treatment, cells were fixed, stained for expression of CDX2, and returned to the microscope for registration with the final time-lapse image. Mean nuclear OCT4-mCherry and CDX2 were quantified for each cell, and the resulting distribution was fit to a 2-component mixed Gaussian distribution representing self-renewing (OCT4+/CDX2-) and differentiated (OCT4-/CDX2+) cells (**Figure EV6**). hESCs that could be assigned to either distribution with >99% confidence (red and green dots, *p_dm_<* 0.01 or *p_dm_>* 0.99) were considered for pro-fate analysis. Cells that did not reach this threshold (gray dots) were not used to determine pro-fate. Because cells in the final frame were assigned to only self-renewing or differentiated categories, pro-mixed cells (yellow traces in panel **A** do not persist to the final frame. **C.** Distributions of OCT4 levels in pro-self-renewing, pro-mixed, and pro-differentiated cell populations. **D.** Distributions of OCT4 burst frequencies in pro-pluripotent, pro-mixed, and pro-differentiated cell populations. **E.** Distributions of cell cycle durations in pro-pluripotent, pro-mixed, and pro-differentiated cell populations. To gain an unbiased look at preexisting determinants of cell fate in panels **D-E**, only cells who completed their entire cell cycle duration before BMP4 addition (t = 0) were included in the analysis. **F.** Correlation between OCT4 level and burst frequency across the entire population of pro-fate cells. **G.** Correlation between OCT4 level and cell cycle duration across the entire population of pro-fate cells. * *P* < 0.05, ** *P* < 0.005, *** *P* < 0.0005; ns, not significant. *ρ*, Pearson correlation; *P, P*-value.

We next asked whether there were preexisting differences between pro-self-renewing, pro-mixed, and pro-differentiated cell populations that might influence their fate decisions. Indeed, pro-self-renewing cells showed significantly higher OCT4 expression levels than either pro-differentiated or pro-mixed populations (**Figure 2C**). This result echoes the observation that repression of Oct4 in mouse ESCs induces loss of pluripotency and differentiation to trophectoderm (Niwa et al., 2000). Pro-self-renewing cells also showed greater burst frequency (number of OCT4 bursts per hour) than pro-mixed cells (**Figure 2D**) and had shorter cell cycle durations than both pro-mixed and pro-differentiated populations (**Figure 2E**). The latter finding is consistent with reports that hESC self-renewal is linked with a shortened G1 cell cycle phase (Becker et al., 2006). Furthermore, we observed that both burst frequency and cell cycle duration were strongly correlated with mean OCT4 levels (**Figure 2F-G**). Taken together, these results show that undifferentiated hESCs display heterogeneous OCT4 levels, burst dynamics, and cell cycle durations. These single-cell features, which were evident as early as 2 days before the differentiation stimulus was presented, were associated with alternate cell fate decisions. Although the preexisting differences between pro-fate populations were statistically significant, these populations of cells still showed considerable heterogeneity and overlapping measurements, suggesting that there are additional factors or events that influence final cell fate.

Because OCT4 levels were the strongest predictors of cell fate (**Figure 2C**), we next asked how heterogeneity in OCT4 levels arises in a population of hESCs. To identify the source of cell-to-cell heterogeneity, we monitored OCT4 expression continuously in proliferating, undifferentiated hESCs for 72 h and generated lineage trees of single-cell relationships (**Figure 3A**). Visual inspection of the lineages revealed that OCT4 levels were most similar among closely related cells (i.e., cells emerging from a common cell division event), providing further support that OCT4 levels are heritable from mother to daughter cell. To quantify this heritability pattern, we calculated the differences in OCT4 levels between pairs of cells as a function of their shared history. Sister cells showed the most similarity in OCT4 levels, followed by “cousin” and “second cousin” cells (**Figure 3B**). Both sister and cousin cells, but not second cousins, were more similar than randomly paired cells, indicating that similarity in OCT4 levels can persist for at least two cell cycle generations (Spencer et al., 2009). Suspecting that each cell division event introduced variability in OCT4 levels, we detected a strong correlation between the number of cell divisions and the difference in OCT4 levels between all pairs of cells (**Figure EV7**). Thus, OCT4 levels are heritable from mother to daughter cell, but each division event introduces incremental variability in OCT4 expression levels.

**Figure 3.**
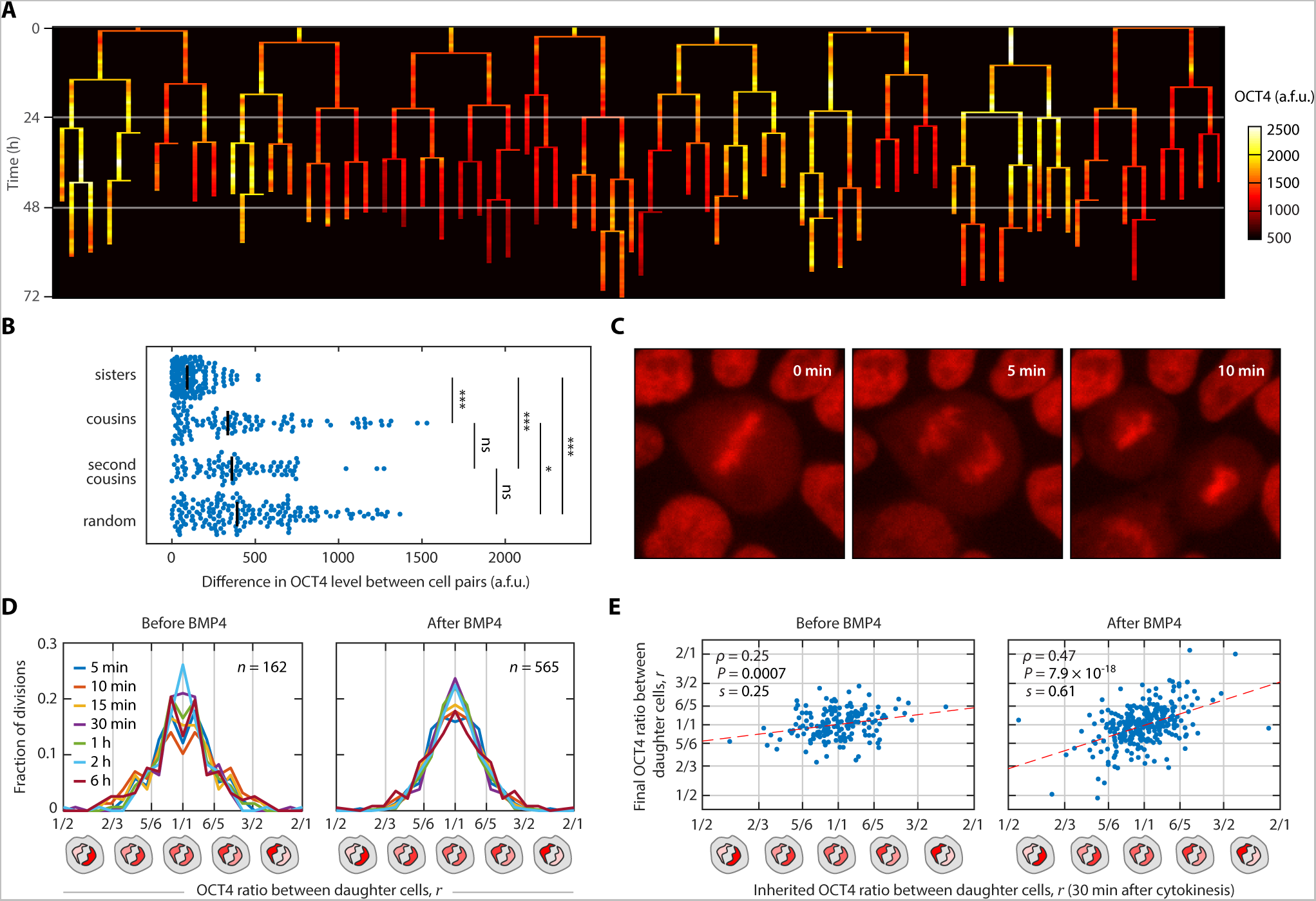
Differences in OCT4 expression levels arise through asymmetric distribution of OCT4 to daughter cells. **A.** Lineage of OCT4 expression dynamics. Mean nuclear OCT4 levels were quantified in individual hESCs continuously for 72 h under undifferentiated conditions. Vertical bars represent individual cells. Thin horizontal bars denote cell division events. Color scale indicates low (*black*), intermediate (*red*), and high (*white*) OCT4 expression levels. **B.** Differences in OCT4 levels between sister cells, cousin cells, second cousin cells, and randomly paired cells. **C.** Filmstrip showing distribution of OCT4 to daughter cells during cell division. **D.** Distribution of OCT4 ratios between sister cells before and after BMP4 treatment. Ratios for both sister cells (*r* and 1/*r*) are plotted to emphasize symmetry. Differently colored curves represent the distribution of ratios at different time points after division. For time points after 5 minutes, the ratio was determined by first calculating a mean OCT4 level for each sister cell among all previous time points and then calculating the resulting ratio between sisters. **E.** Correlation between inherited OCT4 ratio established within 30 minutes of cell division and the final OCT4 ratio between daughters. To avoid trivial correlations, OCT4 measurements used to calculate the initial ratio (x-axis) were excluded from calculation of the final ratio (y-axis). * *P* < 0.05, *** *P* < 0.0005, two-sample Kolmogorov-Smirnov test; ns, not significant. *ρ*, Pearson correlation; *P*, *P*-value; *s*, slope of best fit line.

We then examined cell division events at high temporal resolution to understand how variability in OCT4 levels arises during this process. As cells entered mitosis, OCT4 became visibly associated with the condensed chromosomes (**Figure 3C**, *left panel*). This compacted state persisted throughout anaphase until the two daughter chromatids could be visibly distinguished. We used this first time point—before cytokinesis was complete—to quantify the levels of OCT4 in both newly born daughter cells (**Figure 3C**, *center panel*). Comparison of OCT4-mCherry intensity between daughter cells revealed that the distribution of OCT4 was not perfectly symmetric but instead adopted a bell-shaped distribution that was centered around a mean ratio of 1 (*r* = 1/1) (**Figure 3D**). Approximately 38% of divisions produced daughter cells with *r* = 5/6 or a more extreme ratio; 12% of divisions resulted in *r* = 3/4 or a more extreme ratio; and 3% of division events resulted in *r* = 1/2 or a more extreme ratio. These differences in OCT4 ratios were not due to measurement error because the distribution of *r* between sisters remained consistent for several hours after cytokinesis both before and after BMP4 treatment (see below). In addition, we calculated the half-life of OCT4 to be ~8 h (**Figure EV3**), making it unlikely that asymmetric ratios were due to stochastic differences in protein degradation during the first 5 minutes of daughter cell lifetime (OCT4 would be >99% maternal 5 minutes after division). Moreover, OCT4 ratios were not correlated with nuclear area or radial position within the colony (**Figure EV8**). Thus, significant differences in OCT4 protein levels between sister-cell pairs were established at the moment of cell division.

We next tested whether the ratio of OCT4 inherited by a particular daughter cell influenced its downstream behavior. With a calculated half-life of 7.24 h (**Figure EV3**), the OCT4-mCherry that is established in a daughter cell 30 minutes after division is more than 95% maternal. By comparing OCT4-mCherry intensities between sister chromatids at this time point, we found that the inherited ratio of OCT4 was predictive of the final OCT4 ratio for daughter-cell pairs (**Figure 3E**). Daughter cells receiving the larger proportion of OCT4 (*r* > 1) showed sustained increases in OCT4 relative to the sister cell, whereas daughters receiving the smaller proportion of OCT4 (*r* < 1) showed relative decreases in OCT4. This trend became stronger after BMP4 treatment (**Figure 3E**, *right panel*) and as more time elapsed after cell division (**Figure EV9**). Thus, the ratio of OCT4 established within 30 minutes of cell division is correlated with the amount of OCT4 maintained throughout the lifetime of a cell.

To summarize thus far, differences in OCT4 expression levels arise through asymmetric distribution of OCT4 to daughter cells (**Figures 3C-E**), and these differences are stabilized within the first 10-20 minutes of cellular lifetime (**Figure EV10**). Precise levels of OCT4 are transmitted from mother to daughter cells as reflected by both the similarity among cells that share a common lineage (**Figures 3A-B**) as well as the observation that most progenitor cells (89%) give rise to cellular offspring with the same fate (**Figure 2A**). Taken together with the observation that OCT4 levels are strongly predictive of cell fate decisions (**Figure 2C**), these results suggest that inheritance of OCT4 from the mother cell, including possibly asymmetric distribution during cell division, is strongly determinative of stem cell fate.

To test this idea, we asked at what point during a cell’s lifetime its fate is determined. First, we examined three time periods that were classified as “maternal”, “inherited”, or “autonomous” (**Figure 4A**). Maternal OCT4 encompasses the OCT4 levels in the mother cell up to mitosis (~30 min before daughter cell birth). Inherited OCT4 levels are those established within 30 minutes of cell division, reflecting any asymmetric distribution of OCT4 to daughter cells. Autonomous OCT4 levels include both maternal and inherited levels as well as those experienced during the remainder of the cell’s own lifetime. We considered 304 cells for which we captured their complete cell-cycle lifetime (from birth to division) before BMP4 treatment and for which the pro-fate of self-renewal (286 cells) or differentiation (84 cells) was definitively known. These cells were aligned at the moment of birth to determine the accuracy of predicting cellular fate based on the levels of OCT4 levels in the mother and daughter (**Figure 4B**). At each time point along the mother-daughter timeline, we performed a logistic regression to predict the final fate of the cells as a function of the cumulative OCT4 levels up until that point in time (see **Dataset EV1**, which includes documented code). The logistic regression provides, for each cell, the probability of observing a particular fate given its individual history of OCT4 levels. By comparing predicted probabilities to known fate decisions, we calculated accuracy of the each model based on the percentage of correctly predicted fates (**Figure 4C**).

**Figure 4.**
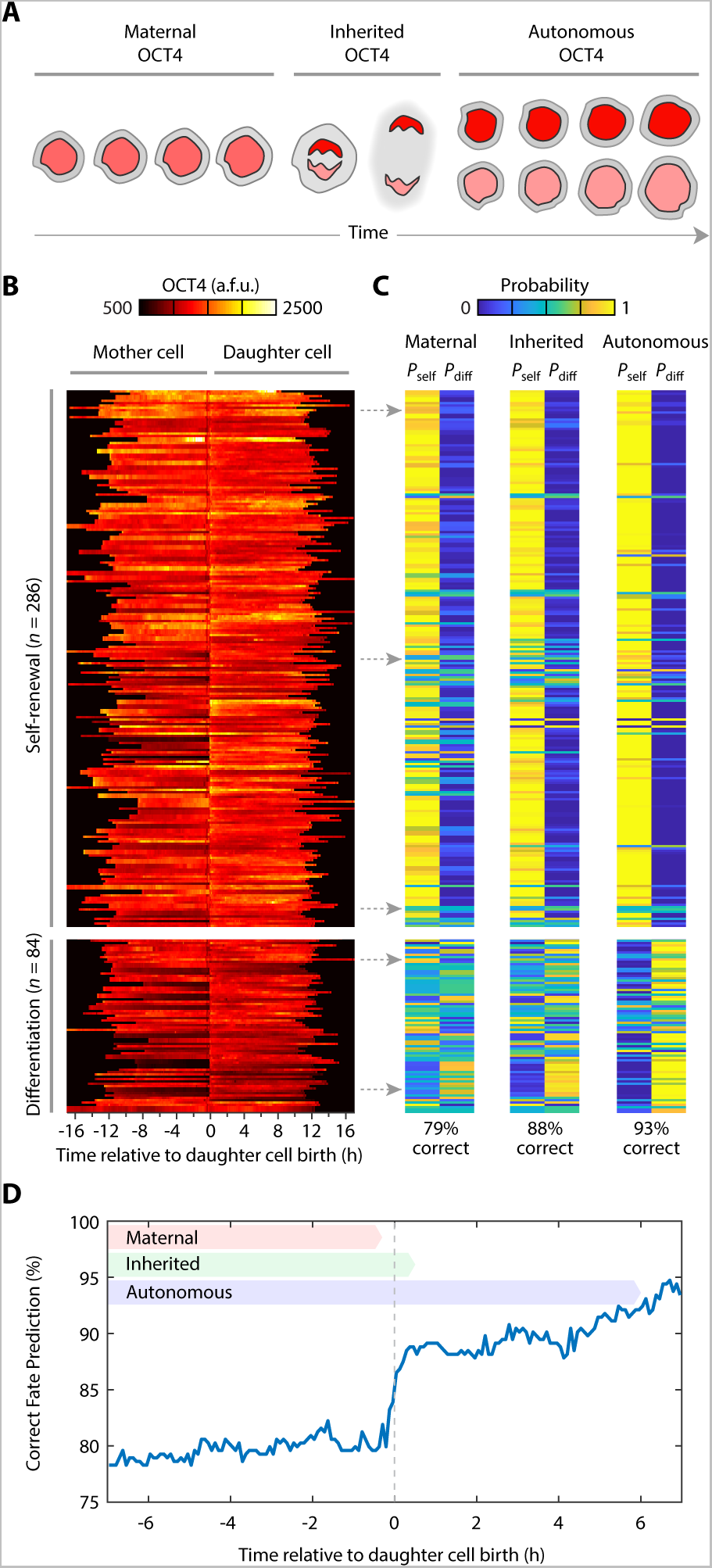
OCT4 levels established during cell division predict fate choice. **A.** Single-cell OCT4 traces were classified into 3 categories: Maternal OCT4 encompasses the levels throughout the lifetime of the mother cell. Inherited OCT4 represents the levels established within 30 minutes of division. Autonomous OCT4 refers to the OCT4 dynamics during the remaining lifetime of the daughter cell. **B.** Single-cell traces of hESCs with either self-renewing or differentiated pro-fates (i.e., all progeny acquiring the same fate) were aligned at the time of daughter-cell birth. Each daughter cell trace (right) was concatenated with its mother-cell trace (left). Color indicates OCT4-mCherry expression levels (reported in a.f.u.) sampled every 5 minutes. **C.** Predicted probabilities for a logistic regression model that determines cell fate based on the history of OCT4-mCherry expression levels. For each cell, the model assigns a probability of being either self-renewing of differentiated based on the history of OCT4 expression dynamics. Each row in the heat maps corresponds to a single mother-daughter cell pair in the same row of panel B. Predicted fate probabilities for maternal OCT4 were calculated by using all OCT4 values up to the moment of division. Probabilities for inherited OCT4 were calculated using values up to 30 minutes after cell birth. Probabilities for autonomous OCT4 were calculated using all OCT4 values up to 6 h after division. **D.** Percentage of correctly predicted fate decisions based on history of OCT4 expression dynamics. At each time point, logistic regression was performed using OCT4 values up to that time point (**Dataset EV1**).

By considering only maternal OCT4 levels before cell birth, the logistic regression model was already 79% correct in predicting cellular fate and rose gradually to 82% over the lifetime of the mother cell (**Figure 4D**). This calculation is consistent with the observation that self-renewing and differentiated cells could be reasonably distinguished by OCT4 signaling in ancestor cells (**Figures 2C-E**). Predictive accuracy improved sharply to 88% within 10 minutes of daughter cell birth, supporting the finding that asymmetric distribution of OCT4 during mitosis has a sudden and lasting influence on daughter cells’ OCT4 levels (**Figure 3E**). During the remaining lifetime of the aligned cells, OCT4 levels became increasingly predictive of cellular fate, reaching 93% within 6 hours of birth. Thus, in this experimental context, differentiation of human embryonic stem cells is almost entirely predetermined both by maternal and mitotic OCT4 expression dynamics, whereas the contribution of autonomous signaling is relatively minor.

In conclusion, we developed an endogenous fluorescent reporter for the canonical pluripotency factor OCT4 to capture differentiation of human embryonic stem cells in real time. We found that the decision to differentiate to embryonic mesoderm is largely determined before the differentiation stimulus is presented to cells and can be predicted by a cell’s preexisting OCT4 levels, bursting frequency, and cell cycle duration. These results in human cells harmonize with studies of mouse ESCs in which cell-to-cell differences in OCT4 expression (Goolam et al., 2016; Niwa et al., 2000; Radzisheuskaya et al., 2013; Zeineddine et al., 2006) and degradation kinetics (Filipczyk et al., 2015; Plachta et al., 2011) are associated with different developmental fate decisions.

However, our results reveal the precise time-window during which significant differences in OCT4 arise. We show that OCT4 levels established immediately after cell division through asymmetric distribution of OCT4 to daughter cells predict its final OCT4 levels. Further, we find that a cell’s fate choice is mostly determined by the time of its birth: hESC differentiation was predicted with ~80% accuracy based on the history of its maternal OCT4 levels alone. Within 30 minutes of birth, this estimate improved to ~90% accuracy. These findings are consistent with a model of mitotic bookmarking in which pluripotency factors such as OCT4 bind tightly to chromatin during mitosis to retain pluripotency gene expression program in daughter cells (Egli et al., 2008; Liu et al., 2017). It is possible that high retention of OCT4 through mitosis essentially seals the fate of daughter cells. If true, this would imply that a cell’s autonomous decision-making may be limited by inherited factors.

This study provides several conceptual advances for the field of single-cell systems biology. First, our work implies that the fate of an individual stem cell is almost entirely determined by the time that cell is born. This idea challenges the view that a cell is a “clean slate” whose fate can be altered by external stimuli. On the contrary, our work suggests that it is more likely the second-or third-generation offspring of a cell that will experience the change in cell fate decision due to external stimuli. As a result, our results call for a reconsideration of the concept of pluripotency—currently defined as the capacity of a given cell to undergo differentiation. Instead, our study argues that pluripotency might be more properly comprehended as a heritable trait that characterizes an entire lineage of proliferating cells. Furthermore, our calculations show that the time period of mitosis is especially important in setting the fate of human embryonic stem cells. There is a significant increase in predictive power within the first few minutes of cell birth that is based on the cell’s level of OCT4. Although a cell retains some autonomy over their fate during their remaining lifetime, a cell’s fate is nearly 90% determined in the moments shortly after it emerges from the mother cell.

## Acknowledgements

We thank Paul Lerou, Galit Lahav, Allon Klein, Jean Cook, Paul Maddox, Bill Marzluff, Scott Bultmann, Peijie Sun, Robert Corty, Greg Keele, and members of the Purvis Lab for helpful discussions and technical suggestions. This work was supported by NIH grant DP2-HD091800-01, the W.M. Keck Foundation, and the Loken Stem Cell Fund.

## Author Contributions

S.C.W., R.D., and J.C. constructed the OCT4-mCherry reporter cell line. S.C.W. and R.D. performed validation studies. S.C.W. and C.D. performed live-cell imaging. C.D., R.A.H., T.M.Z. and K. K. conducted image analysis and cell tracking. C.D., R.A.H., and J.E.P. performed computational analysis. S.C.W. and A.S.B. carried out lineage differentiation experiments. J.E.P wrote the manuscript with contributions from all authors.

## Materials and Methods

### Culture and treatment of hESCs

WA09 (H9) hES cell line was purchased from WiCell (Wisconsin) and maintained in mTeSR1 (05850, StemCell Technologies) on growth factor reduced Matrigel (354230, BD). Cells were passaged every three days using 0.5% EDTA in PBS.

### Guide RNA and CRISPR/Cas9 cutting vector

The gRNA sequence GTGAAATGAGGGCTTGCGA, targeting the start codon of human *POU5F1* (OCT4), was cloned into pX330 (AddGene) using the standard cloning protocol described Ran et al(Ran et al., 2013). The cutting efficiency of the Cas9/OCT4-gRNA was validated with Guide-it Mutation Detection Kit (Takara Bio).

### Donor cassette construction

The 5’ homology arm of OCT4 was amplified out of H9 genomic DNA with the following primers (Fwd: 5’-AAGGTTGGGAAACTGAGGCC-3’, Rev: 5’-GGGAAGGAAGGCGCCCCAAG-3) yielding a 1114 bp homology arm that was then cloned into the pGEMTEZ plasmid (Promega) followed by the coding sequence for the mCherry fluorescent protein (minus its stop codon) followed by a short linker sequence (TCC GGA TCC) and the start ATG codon for OCT4. The OCT4 gene constituted the 3’ homology arm and was amplified out of H9 genomic DNA with the following primers (Fwd: 5’- ATGGCGGGACACCTGGCTTC-3’, Rev: 5- AGCTTTCTACAAGGGGTGCC-3’) yielding a 1082 bp homology arm.

### Introduction of exogenous DNA into H9 cells

H9 cells were cultured on 10 cm dishes and, when 80% confluent, were dissociated using 0.5mM EDTA. 10 × 10^6^ cells were resuspended in 800 μL ice-cold PBS containing 25 μg of the OCT4-mCherry donor vector and 25 μg of the guideRNA/Cas9 vector. Cells were electroporated in 100 μL tips (Neon, ThermoFisher Scientific) using program 19 of the optimization protocol (1050V, 30ms, 2 pulses) and resuspended in mTeSR1 (STEMCELL Technologies) supplemented with Rock inhibitor (S1049, Selleck Chemicals) at a final concentration of 10 μM. When colonies that expressed mCherry reached approximately 20 mm in size, they were marked and picked into Matrigel coated 24-well plates.

### Endogenous OCT4 levels

Endogenous OCT4 levels in H9 wild-type cells and H9 OCT4-mCherry clone 8-2 were determined by antibody staining using a rabbit anti-OCT4 antibody (ab19857, Abcam). Immunostaining was performed using standard protocols. Briefly, cells were fixed for 15 min in 4% paraformaldehyde and permeabilized and blocked for 30 minutes in 5% goat serum with 0.3% Triton X-100 in TBS. Incubation with primary antibody was performed overnight and the incubation with the secondary antibody (Molecular Probes) was done at room temperature for 45 minutes. Nuclei were visualized using NucBlue Fixed Cell Stain ready Probes reagent (R37606, Molecular Probes).

### Live-cell imaging

Asynchronous H9 OCT4-mCherry cells were plated on 12-well glass bottom plates (Cellvis) in phenol-red free or clear DMEM/F-12 (Gibco) supplemented with mTeSR1 supplement (05850, STEMCELL Technologies) approximately 24 hours before being imaged. Cells were imaged using a Nikon Ti Eclipse microscope operated by NIS Elements software V4.30.02 with an Andor ZYLA 4.2 cMOS camera and a custom stage enclosure (Okolabs) to ensure constant temperature, humidity, and CO2 levels. Fresh media with or without BMP4 was added every 24 h. Images were flat-field corrected using NIS Elements.

### Image analysis

A custom ImageJ plugin (available upon request) was used to perform automated segmentation and manually tracking of hESCs. Fluorescence intensity was quantified using an adapted threshold followed by watershed segmentation of the OCT4-mCherry channel. The program tracked the cell ID, parent ID, frame number, mean intensity and exported this information to MATLAB for analysis.

### Quantitative analysis

All computational methods including lineage analysis and logistic regression are included as **Dataset EV**, which includes processed image data and documented MATLAB code used to generate each of the figures.

---

**Figure EV1.**
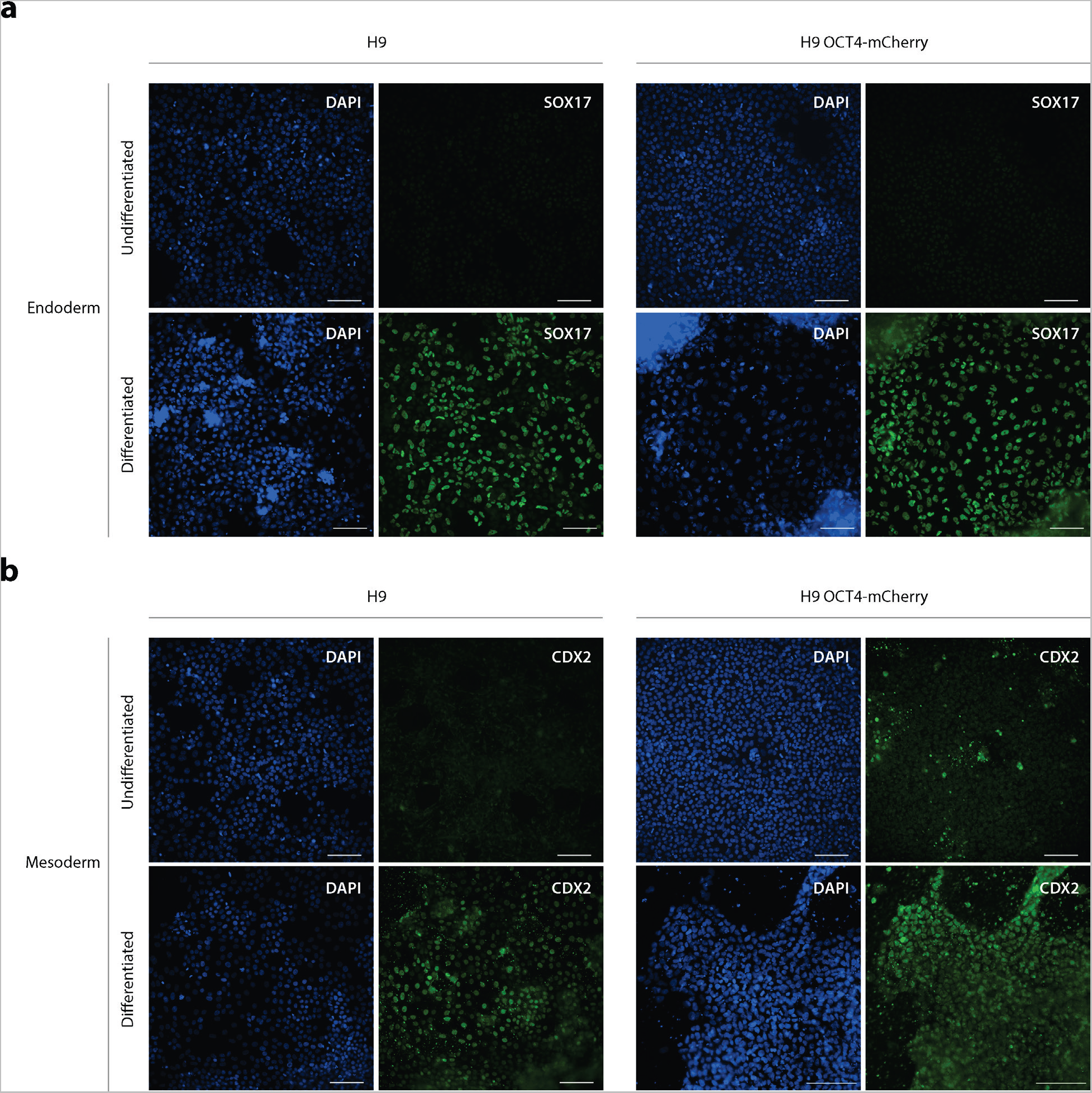
hESCs with high OCT4 expression after BMP4 treatment are competent to differentiate into multiple cell types. H9 wild-type and H9 OCT4-mCherry cells were seeded at a density of 2 × 10^5^ cells/cm^2^. After 1 d, cells were treated with 100 ng/ml BMP4 (Peprotech). After 24 hours, the center “core” of each colony (see **Figure 2**) was manually removed and placed on Matrigel-coated 24-well glass bottom plates. Cells were allowed to adhere for 2 hours and then differentiated toward **a**, endoderm or **b**, mesoderm using the STEMdiff™ Trilineage Differentiation Kit (STEMCELL Technologies). Scale bar = 100 μm.

**Figure EV2.**
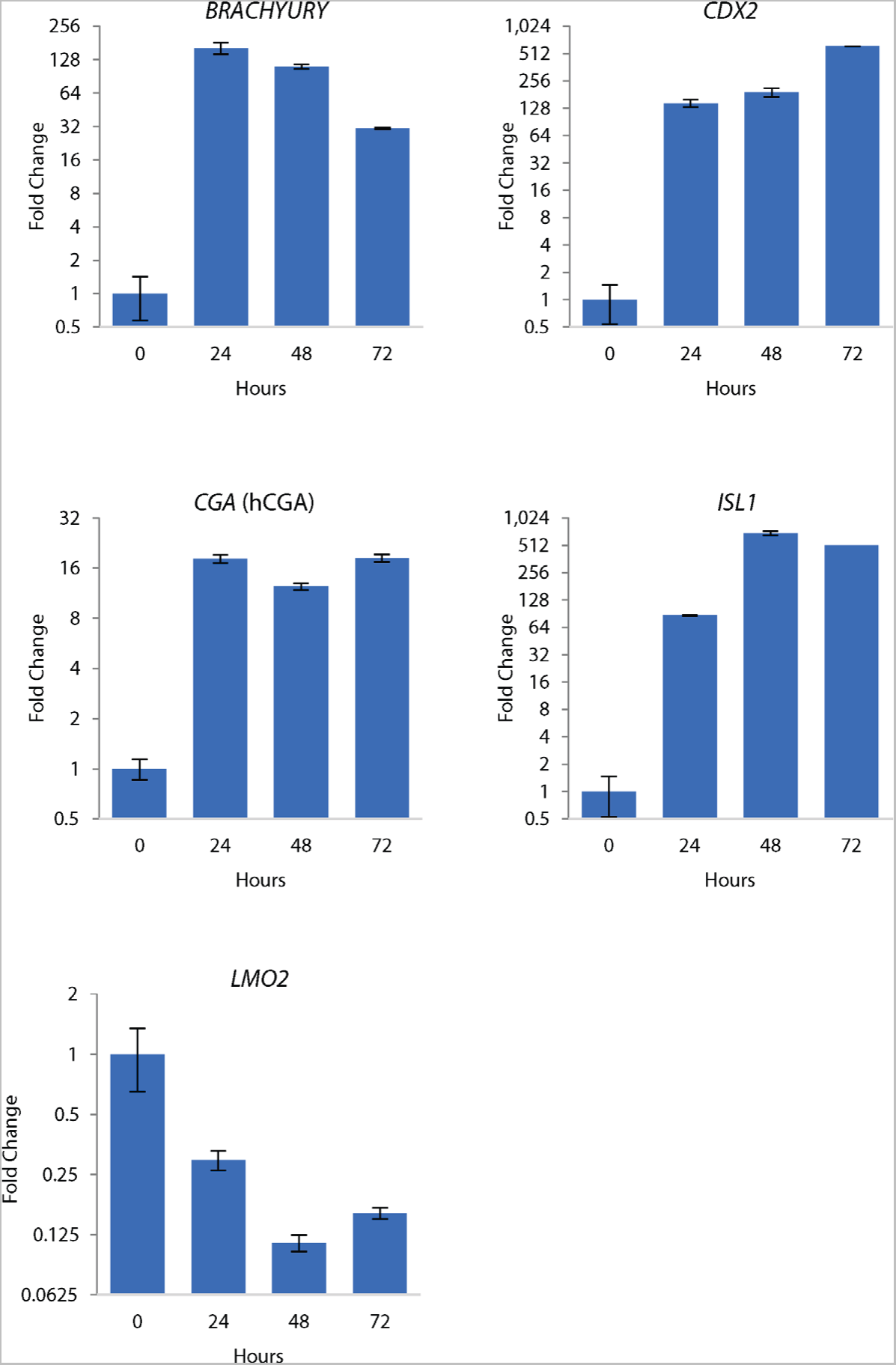
Expression of mesoderm-specific markers in response to BMP4 treatment. H9 hESCs were treated with 100 ng/mL BMP4 and harvested at 24, 48, and 72 h for quantification of transcript levels.

**Figure EV3.**
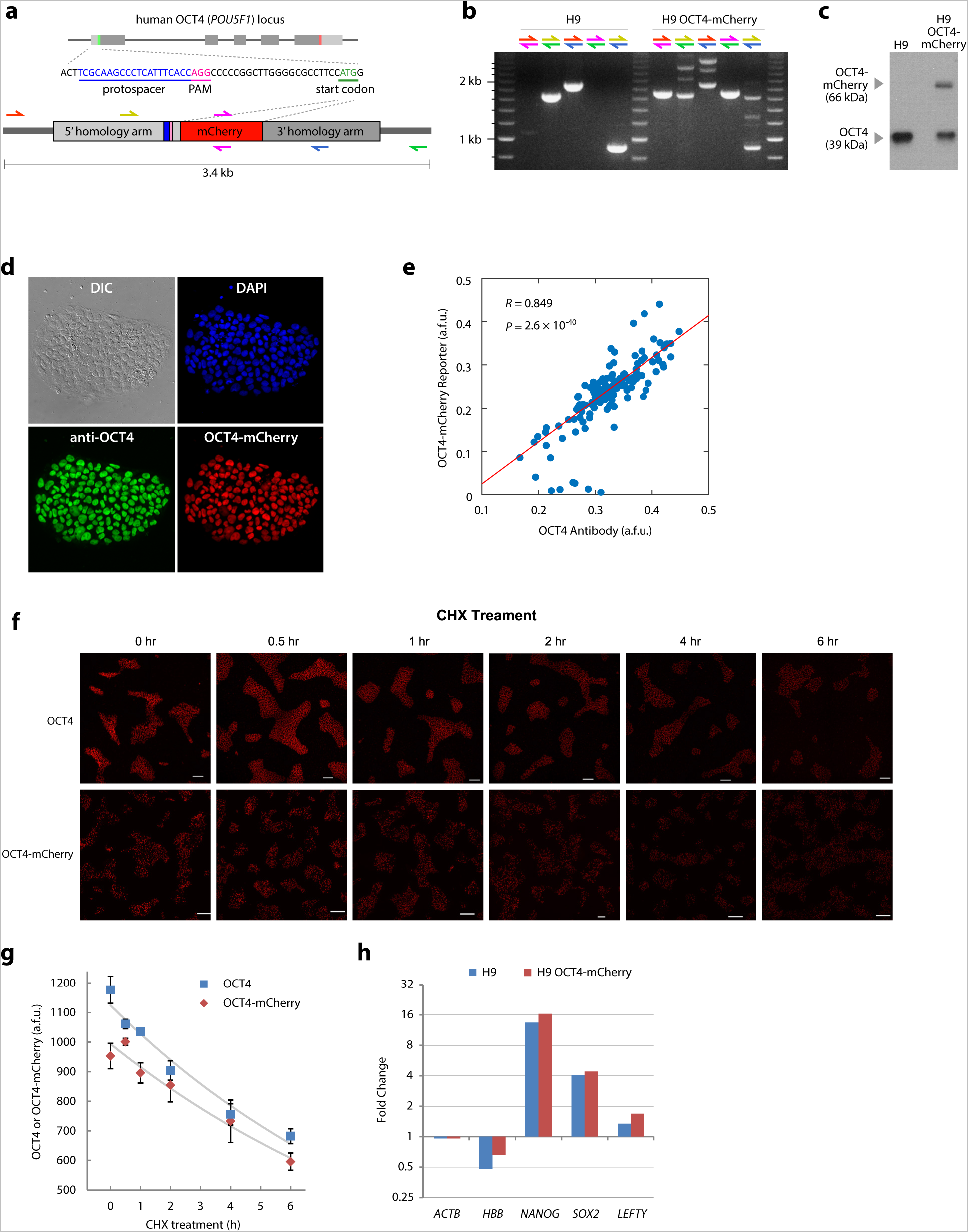

**Figure EV4.**
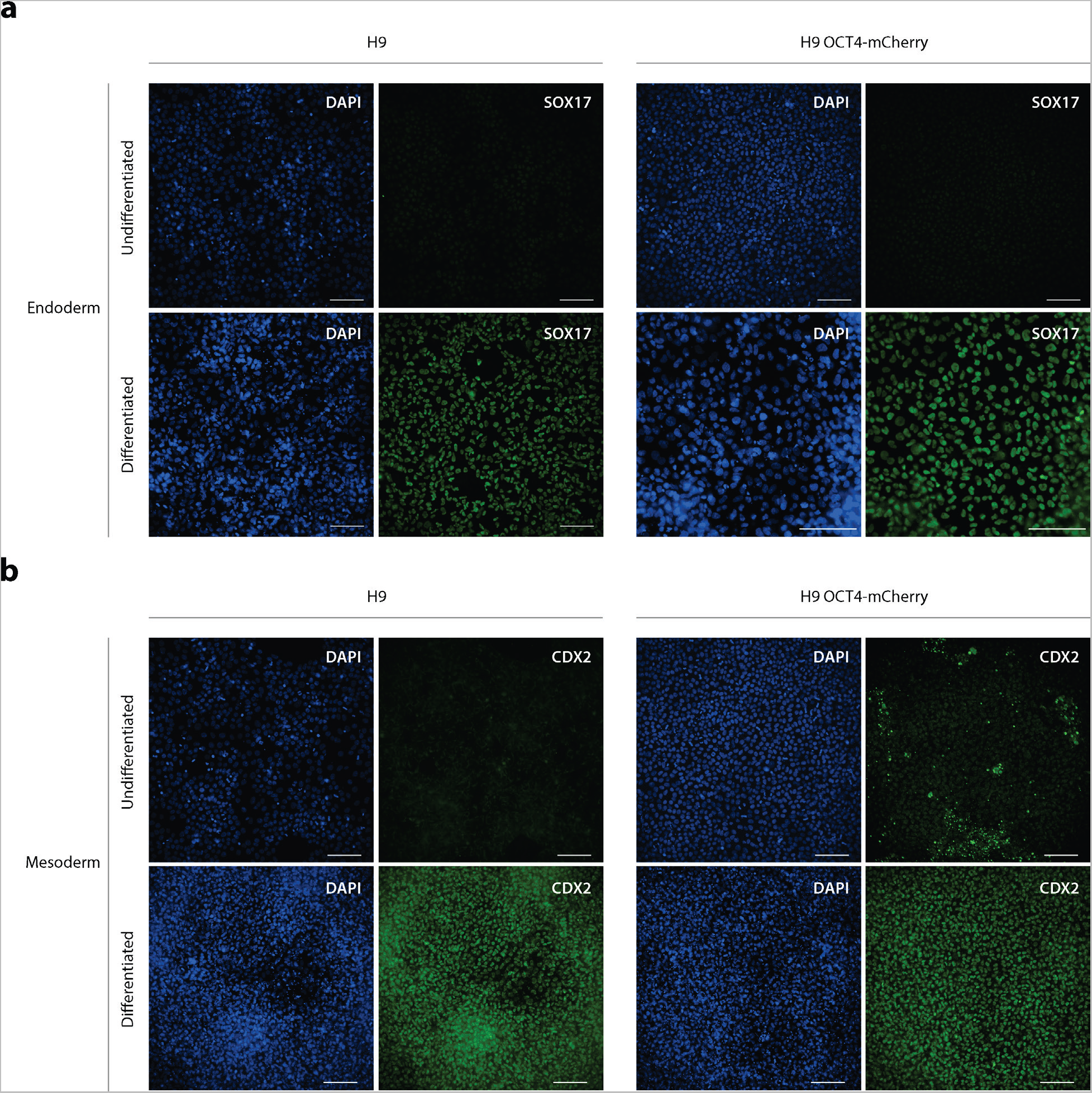
Cells bearing the OCT4-mCherry reporter are competent to differentiate into multiple cell types. H9 wild-type and H9 OCT4-mCherry clone 8-2 cells were seeded at the density of 2 × 10^5^ cell/cm^2^ for endoderm and 5 10^4^ cell/cm^2^ for mesoderm lineage differentiation on 24-well glass bottom plates using mTeRS1 medium supplemented with 5 μM Y27632 (STEMCELL Technologies). After 24 hours, cells were differentiated into **a**, endoderm or **b**, mesoderm using the STEMdiff™ Trilineage Differentiation kit (STEMCELL Technologies). Medium was replaced every day for 5 days, then cells were fixed 15 min with 4% paraformaldehyde, permeabilized for 15 min with 0.3% Triton X-100 and blocked 1 hour with 5% BSA at room temperature. Then, cells differentiated to endoderm were incubated with SOX17

**Figure EV5.**
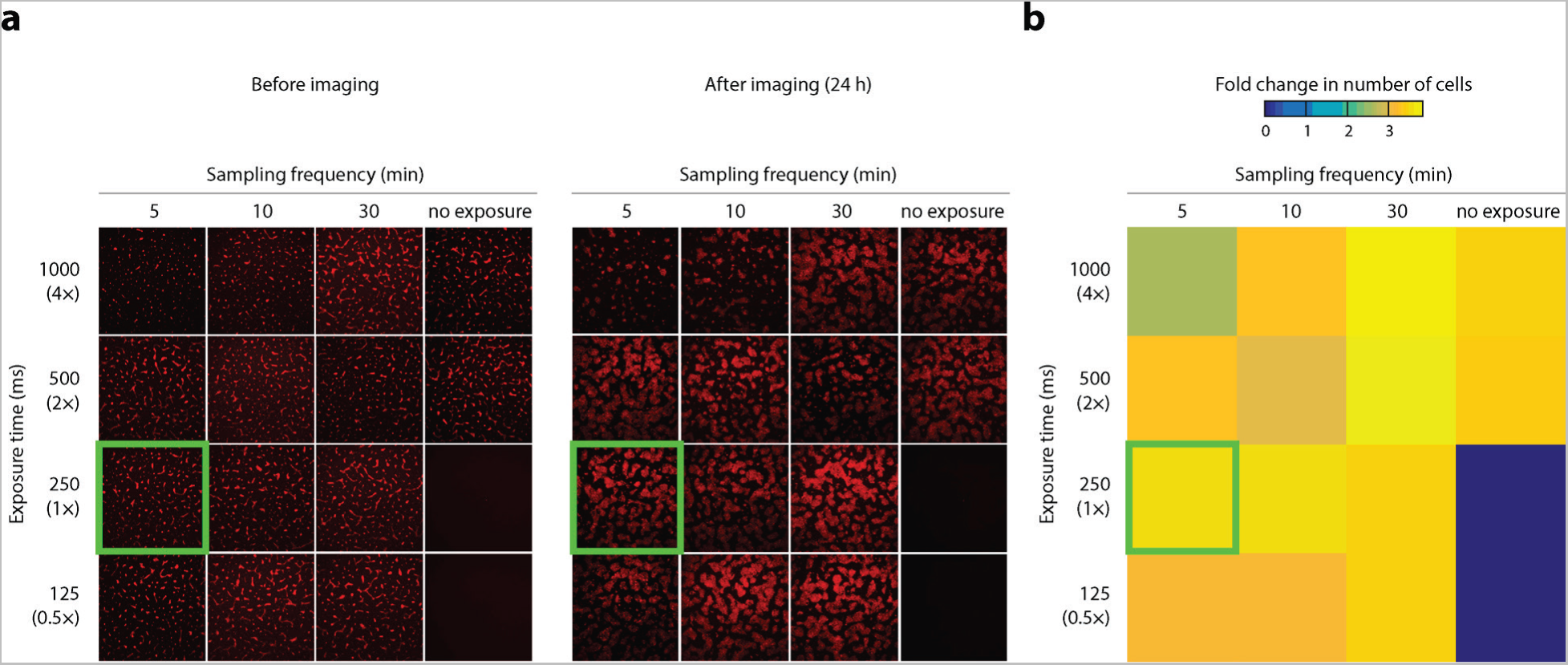
Proliferation of hESCs as a function of light exposure sampling rate and intensity. **a**, H9 OCT4-mCherry cells were seeded onto glass bottom dishes and subjected to time-lapse fluorescence imaging for 24 h at the indicated exposure durations and sampling frequencies. For each condition, a single image was captured by the camera both before and after light exposure to determine the extent of cell proliferation. Nikon Elements spot detection algorithm was used to count nuclei before and after imaging for each condition. In the fourth column, cells were not exposed to any light other than during the first and last image captured for quantification of cell proliferation. The green box indicates the exposure setting used in this study. **b**, Heat map showing the fold change in the number of nuclei quantified for each condition.

**Figure EV6.**
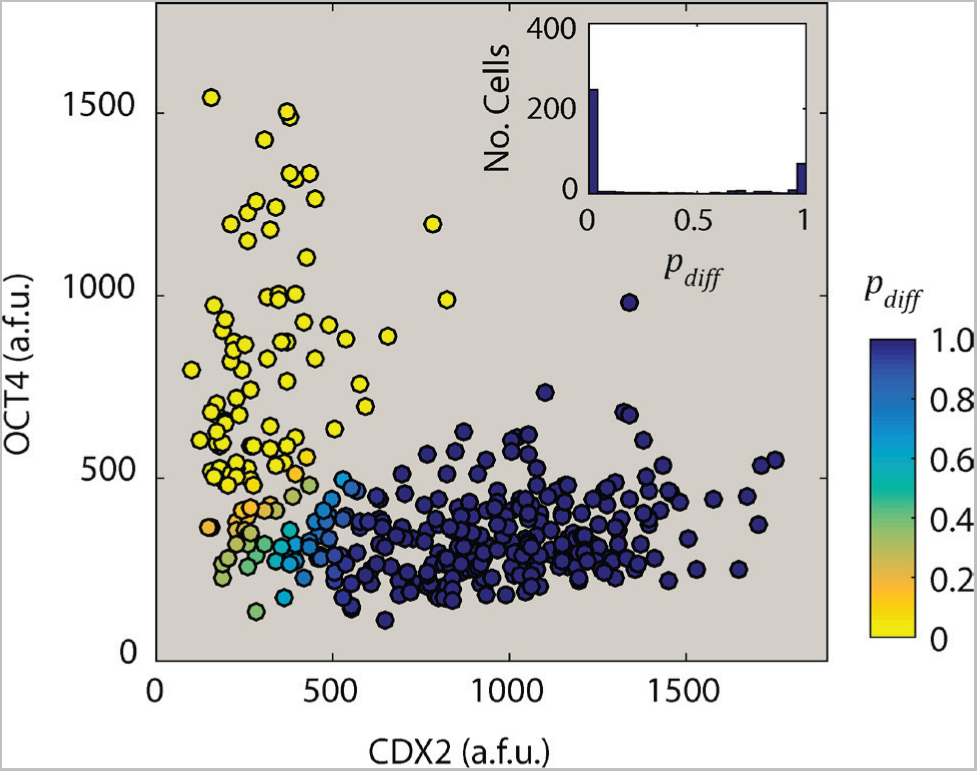
Classification of cell fates using a 2-component Gaussian distribution. Mean nuclear intensity values for OCT4 and CDX2 were used to separate the population of cells into two groups using a mixed Gaussian model. *Inset*, distribution of posterior probabilities (*p*_*diff*_) for all cells. Only cells that could be confidently labeled as pluripotent (*p*_*diff*_<0.01) or differentiated (*p*_*diff*_> 0.99) were considered for pro-fate analysis. Similar class assignments were made by employing *k*-means clustering with *k* = 2.

**Figure EV7.**
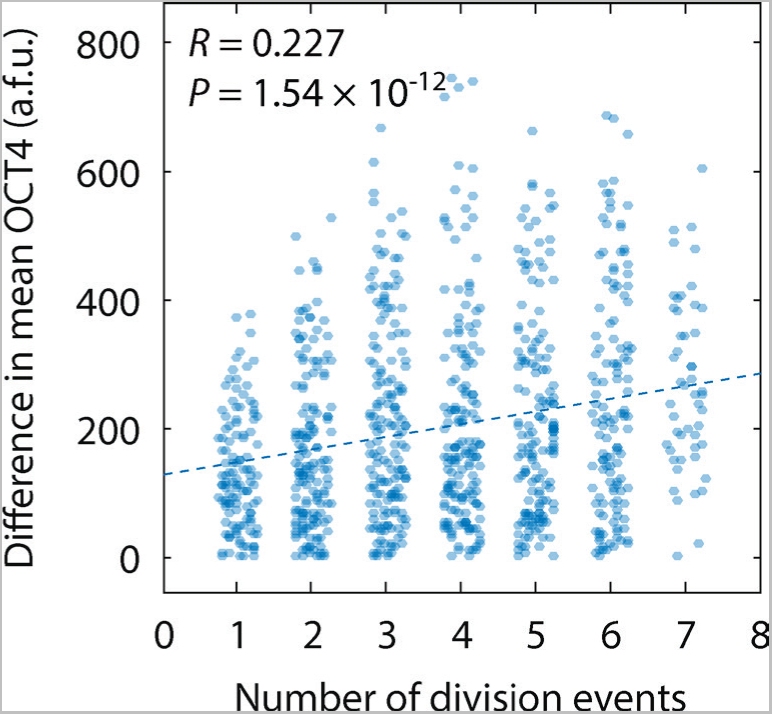
Difference in OCT4 levels is strongly correlated with number of cell divisions separating two cells. All pairs of cells in Figure 3a from the main text were compared for difference in OCT4 level (*y-axis*) and number of cell division events separating those cells (*x-axis*). For example, mother and daughter cell pairs are separated by 1 cell division event, whereas cousin cells are separated by 3 division events. As the number of division events increases, the difference in OCT4 levels off because each division event is equally likely to increase or decrease the difference in OCT4 levels between two given cells.

**Figure EV8.**
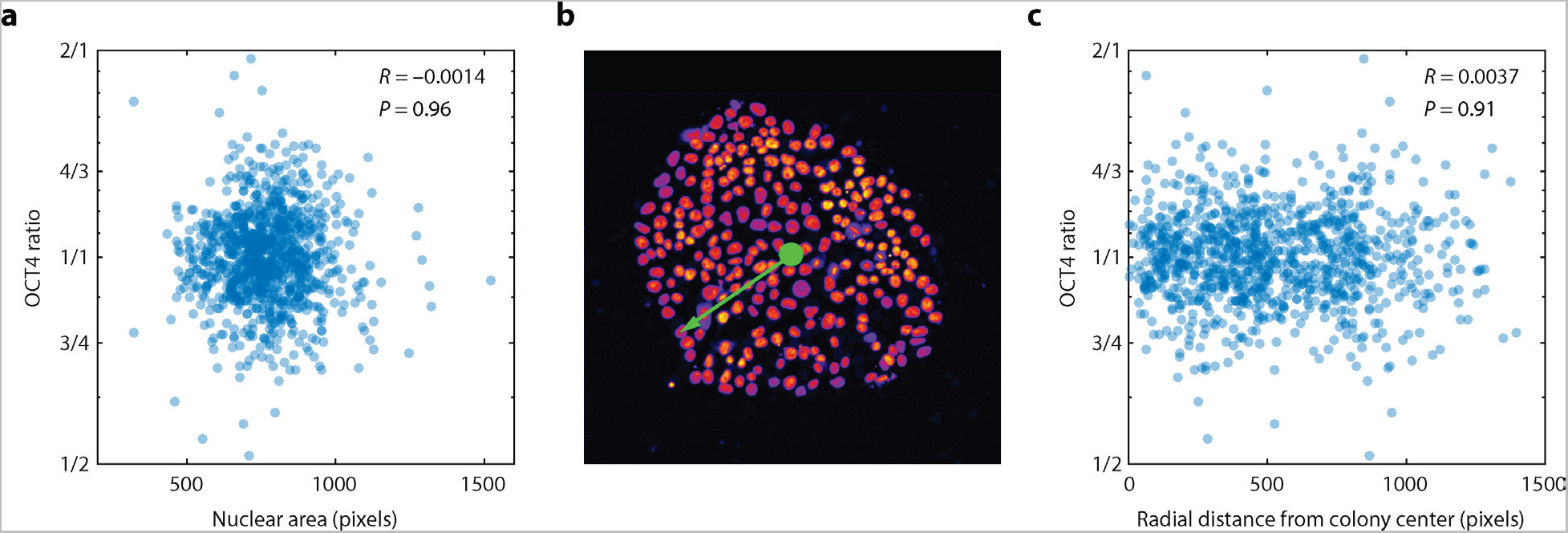
OCT4 ratios are not correlated with nuclear area or radial position within the colony. **a**, Scatter plot of OCT4 ratio as a function of nuclear area for sister cells. To avoid noise due to segmentation errors, nuclear area was calculated 25 min after cell division, when OCT4 became diffuse throughout the nucleus.**b**, The colony center was defined as a center of mass of all OCT4-expressing cells. The radial position of individual cells was defined as a distance (in pixels) between the center of a cell and the center of the colony. **c**, Scatter plot of OCT4 ratio as a function of radial distance from the center of the colony.

**Figure EV9.**
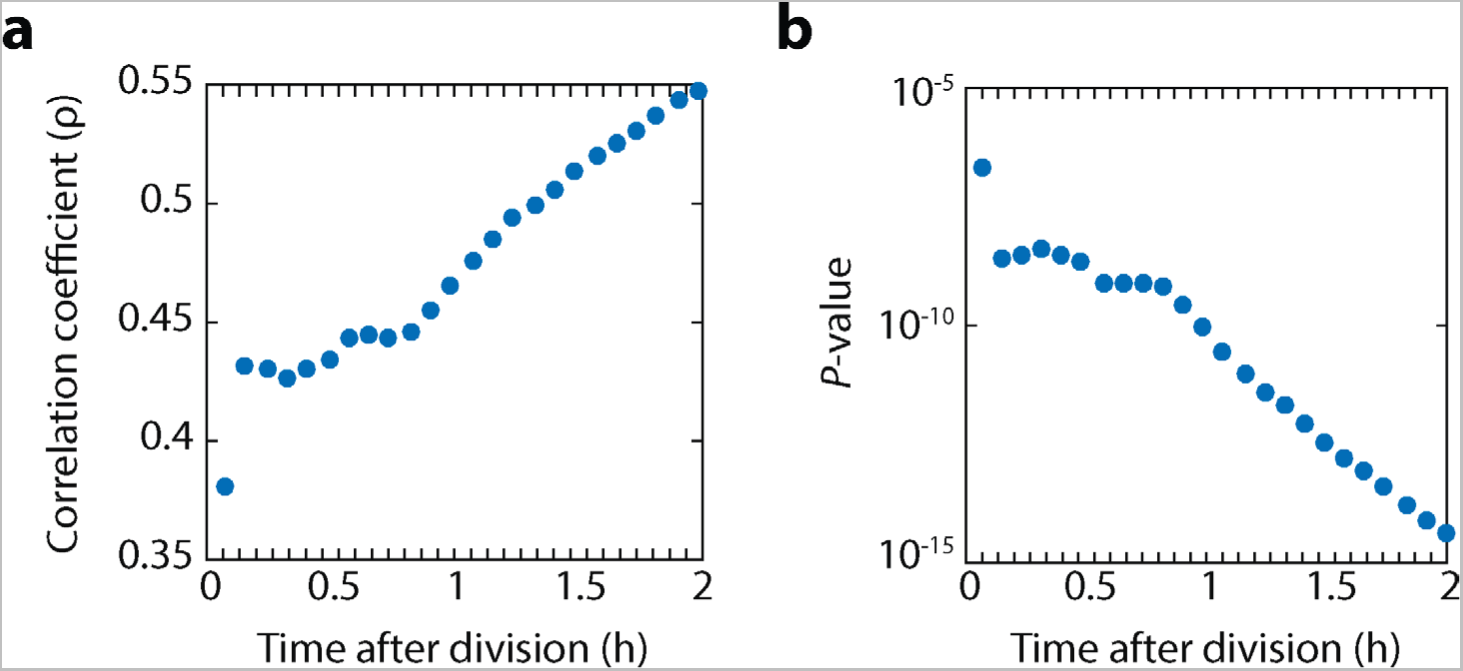
The inherited ratio of OCT4 becomes more strongly correlated with final OCT4 level over time. **a**, Correlation of OCT4 ratio between sister cells and differences in final OCT4 levels. At each time point, the ratio of OCT4 between sister cell pairs was calculated as the ratio of the cumulative sum of mean OCT4 intensity values at all preceding time points. This ratio was used to predict the final OCT4 level for each cell, calculated as the cumulative sum of mean intensities for the remaining time points. **b**, Corresponding *P*-values for correlation in panel **a**.

## References

Albeck, J.G., Mills, G.B., and Brugge, J.S. (2013). Frequency-Modulated Pulses of ERK Activity Transmit Quantitative Proliferation Signals. Molecular cell 49, 249–261.

Arora, M., Moser, J., Phadke, H., Basha, A.A., and Spencer, S.L. (2017). Endogenous Replication Stress in Mother Cells Leads to Quiescence of Daughter Cells. Cell Rep 19, 1351–1364.

Barr, A.R., Cooper, S., Heldt, F.S., Butera, F., Stoy, H., Mansfeld, J., Novak, B., and Bakal, C. (2017). DNA damage during S-phase mediates the proliferation-quiescence decision in the subsequent G1 via p21 expression. Nat Commun 8, 14728.

Becker, K.A., Ghule, P.N., Therrien, J.A., Lian, J.B., Stein, J.L., van Wijnen, A.J., and Stein, G.S. (2006). Self-renewal of human embryonic stem cells is supported by a shortened G1 cell cycle phase. J Cell Physiol 209, 883–893.

Bernardo, A.S., Faial, T., Gardner, L., Niakan, K.K., Ortmann, D., Senner, C.E., Callery, E.M., Trotter, M.W., Hemberger, M., Smith, J.C., et al. (2011). BRACHYURY and CDX2 mediate BMP-induced differentiation of human and mouse pluripotent stem cells into embryonic and extraembryonic lineages. Cell Stem Cell 9, 144–155.

Deglincerti, A., Croft, G.F., Pietila, L.N., Zernicka-Goetz, M., Siggia, E.D., and Brivanlou, A.H. (2016). Self-organization of the in vitro attached human embryo. Nature 533, 251–254.

Egli, D., Birkhoff, G., and Eggan, K. (2008). Mediators of reprogramming: transcription factors and transitions through mitosis. Nature reviews 9, 505–516.

Etoc, F., Metzger, J., Ruzo, A., Kirst, C., Yoney, A., Ozair, M.Z., Brivanlou, A.H., and Siggia, E.D. (2016). A Balance between Secreted Inhibitors and Edge Sensing Controls Gastruloid Self-Organization. Developmental cell 39, 302–315.

Filipczyk, A., Marr, C., Hastreiter, S., Feigelman, J., Schwarzfischer, M., Hoppe, P.S., Loeffler, D., Kokkaliaris, K.D., Endele, M., Schauberger, B., et al. (2015). Network plasticity of pluripotency transcription factors in embryonic stem cells. Nat Cell Biol 17, 1235–1246.

Ghule, P.N., Medina, R., Lengner, C.J., Mandeville, M., Qiao, M., Dominski, Z., Lian, J.B., Stein, J.L., van Wijnen, A.J., and Stein, G.S. (2011). Reprogramming the pluripotent cell cycle: restoration of an abbreviated G1 phase in human induced pluripotent stem (iPS) cells. J Cell Physiol 226, 1149–1156.

Goolam, M., Scialdone, A., Graham, S.J., Macaulay, I.C., Jedrusik, A., Hupalowska, A., Voet, T., Marioni, J.C., and Zernicka-Goetz, M. (2016). Heterogeneity in Oct4 and Sox2 Targets Biases Cell Fate in 4-Cell Mouse Embryos. Cell 165, 61–74.

Lane, K., Van Valen, D., DeFelice, M.M., Macklin, D.N., Kudo, T., Jaimovich, A., Carr, A., Meyer, T., Pe’er, D., Boutet, S.C., et al. (2017). Measuring Signaling and RNA-Seq in the Same Cell Links Gene Expression to Dynamic Patterns of NF-kappaB Activation. Cell Syst 4, 458–469 e455.

Lin, Y., Sohn, C.H., Dalal, C.K., Cai, L., and Elowitz, M.B. (2015). Combinatorial gene regulation by modulation of relative pulse timing. Nature 527, 54–58.

Liu, Y., Pelham-Webb, B., Di Giammartino, D.C., Li, J., Kim, D., Kita, K., Saiz, N., Garg, V., Doane, A., Giannakakou, P., et al. (2017). Widespread Mitotic Bookmarking by Histone Marks and Transcription Factors in Pluripotent Stem Cells. Cell Rep 19, 1283–1293.

Morrison, S.J., and Kimble, J. (2006). Asymmetric and symmetric stem-cell divisions in development and cancer. Nature 441, 1068–1074.

Nemashkalo, A., Ruzo, A., Heemskerk, I., and Warmflash, A. (2017). Morphogen and community effects determine cell fates in response to BMP4 signaling in human embryonic stem cells. Development 144, 3042–3053.

Nichols, J., Zevnik, B., Anastassiadis, K., Niwa, H., Klewe-Nebenius, D., Chambers, I., Scholer, H., and Smith, A. (1998). Formation of pluripotent stem cells in the mammalian embryo depends on the POU transcription factor Oct4. Cell 95, 379–391.

Niwa, H., Miyazaki, J., and Smith, A.G. (2000). Quantitative expression of Oct-3/4 defines differentiation, dedifferentiation or self-renewal of ES cells. Nat Genet 24, 372–376.

Plachta, N., Bollenbach, T., Pease, S., Fraser, S.E., and Pantazis, P. (2011). Oct4 kinetics predict cell lineage patterning in the early mammalian embryo. Nat Cell Biol 13, 117–123.

Purvis, J.E., Karhohs, K.W., Mock, C., Batchelor, E., Loewer, A., and Lahav, G. (2012). p53 dynamics control cell fate. Science 336, 1440–1444.

Radzisheuskaya, A., Chia Gle, B., dos Santos, R.L., Theunissen, T.W., Castro, L.F., Nichols, J., and Silva, J.C. (2013). A defined Oct4 level governs cell state transitions of pluripotency entry and differentiation into all embryonic lineages. Nat Cell Biol 15, 579–590.

Ran, F.A., Hsu, P.D., Wright, J., Agarwala, V., Scott, D.A., and Zhang, F. (2013). Genome engineering using the CRISPR-Cas9 system. Nature protocols 8, 2281–2308.

Spencer, S.L., Gaudet, S., Albeck, J.G., Burke, J.M., and Sorger, P.K. (2009). Non-genetic origins of cell-to-cell variability in TRAIL-induced apoptosis. Nature 459, 428–432.

Suda, J., Suda, T., and Ogawa, M. (1984a). Analysis of differentiation of mouse hemopoietic stem cells in culture by sequential replating of paired progenitors. Blood 64, 393–399.

Suda, T., Suda, J., and Ogawa, M. (1984b). Disparate differentiation in mouse hemopoietic colonies derived from paired progenitors. Proceedings of the National Academy of Sciences of the United States of America 81, 2520–2524.

Suel, G.M., Garcia-Ojalvo, J., Liberman, L.M., and Elowitz, M.B. (2006). An excitable gene regulatory circuit induces transient cellular differentiation. Nature 440, 545–550.

Symmons, O., and Raj, A. (2016). What’s Luck Got to Do with It: Single Cells, Multiple Fates, and Biological Nondeterminism. Molecular cell 62, 788–802.

Thomson, J.A., Itskovitz-Eldor, J., Shapiro, S.S., Waknitz, M.A., Swiergiel, J.J., Marshall, V.S., and Jones, J.M. (1998). Embryonic stem cell lines derived from human blastocysts. Science 282, 1145–1147.

Xu, R.H., Chen, X., Li, D.S., Li, R., Addicks, G.C., Glennon, C., Zwaka, T.P., and Thomson, J.A. (2002). BMP4 initiates human embryonic stem cell differentiation to trophoblast. Nature biotechnology 20, 1261–1264.

Yang, H.W., Chung, M., Kudo, T., and Meyer, T. (2017). Competing memories of mitogen and p53 signalling control cell-cycle entry. Nature 549, 404–408.

Zeineddine, D., Papadimou, E., Chebli, K., Gineste, M., Liu, J., Grey, C., Thurig, S., Behfar, A., Wallace, V.A., Skerjanc, I.S., et al. (2006). Oct-3/4 dose dependently regulates specification of embryonic stem cells toward a cardiac lineage and early heart development. Developmental cell 11, 535–546.

